# Myosin-1b interacts with UNC45A and controls intestinal epithelial morphogenesis

**DOI:** 10.1101/2021.09.09.459609

**Authors:** Céline Revenu, Marianna Parlato, Marion Rosello, Karine Duroure, Rémi Duclaux-Loras, Ophélie Nicolle, Marie-Thérèse Prospéri, Julie Stoufflet, Juliette Vougny, Corinne Lebreton, Priscilla Lépine, Grégoire Michaux, Nadine Cerf-Benssusan, Evelyne Coudrier, Filippo Del Bene

## Abstract

Vesicle trafficking and the establishment of apico-basal polarity are essential processes in epithelium morphogenesis. Myosin-1b, an actin-motor able to bind membranes, regulates membrane shaping and vesicle trafficking. Here, we investigate Myosin-1b function in gut morphogenesis and congenital disorders using cell line and zebrafish larvae as well as patient biopsies. In a 3D Caco-2 cyst model, lumen formation is impaired in absence of Myosin-1b. In zebrafish, both Morpholino knock-down and genetic mutation of *myo1b* result in intestinal bulb epithelium folding defects associated with vesicle accumulation, reminiscent of a villous atrophy phenotype. We show that Myosin-1b interacts with the chaperone UNC45A, genetic deletion of which also results in gut folding defects in zebrafish. Loss of function mutations in *UNC45A* have been reported in complex hereditary syndromes, notably exhibiting intestinal disorders associated with villous atrophy. In UNC45A-depleted cells and in patient biopsies, Myosin-1b protein level is strikingly decreased. The appearance of Myosin-1b aggregates upon proteasome inhibition in cells points at a degradation mechanism of misfolded Myosin-1b in the absence of its chaperone. In conclusion, Myosin-1b plays an unexpected role in the development of the intestinal epithelium folds or villi downstream UNC45A, establishing its role in the gut defects reported in UNC45A patients.

**Summary statement:** Myosin-1b is important for intestinal epithelium folding during zebrafish development and participates in the villous atrophy clinical manifestation downstream UNC45A loss of function.

## Introduction

The establishment of apico-basal polarity and lumen formation are two fundamental steps during vertebrate intestinal epithelial morphogenesis (Chin et al., 2017). The actin cytoskeleton and the vectorial vesicle trafficking play a major role in the initiation and maintenance of this process, leading to a stable single layer of cells with distinct apical and basolateral domains (Lubarsky and Krasnow, 2003, Martin-Belmonte and Mostov, 2008). The apical membrane of the enterocyte is further organized in microvilli, plasma membrane protrusions, which are supported by bundles of parallel actin filaments and interacting proteins interconnected at the basis through a network of actin, spectrin, and myosins known as terminal web (Revenu et al., 2004). The interaction between neighboring polarized cells is further strengthened by the formation of cadherin-based adherens junctions and claudin-based tight junctions. A proper polarization of the intestinal epithelium is essential to achieve its main physiological roles, such as fluids and nutrient absorption and secretion. Indeed, defects in intestinal epithelial cell polarity and apical lumen formation result in early onset intestinal disorders, usually appearing in the first days of life (Kwon et al., 2020). Recently, loss of function (LOF) mutations in the chaperone *UNC45A* were reported in families presenting complex phenotypes including congenital diarrhea and several degrees of villous atrophy (Esteve et al., 2018). UNC45A belongs to the conserved UCS protein family (UNC-45/CRO1/She4p) of myosin co-chaperones.

Myosins 1 are single-headed actin motors targeted to membranes. Myosin1b (Myo1b) was detected in mouse enterocyte brush borders in a mass spectrometry analysis (Revenu et al., 2012). Studies in cell cultures reported that Myo1b associates with organelles and regulates membrane trafficking by controlling their morphology (Almeida et al., 2011). Myo1b can pull out membrane tubes along actin bundles immobilized on a solid substrate (Yamada et al., 2014) and it controls the formation of repulsive filopodia, the redistribution of actomyosin fibres driving cell repulsion (Prosperi et al., 2015) as well as the formation of axons in cultured neurons by controlling actin waves (Iuliano et al., 2018). Despite this progress in understanding Myo1b function *in vitro* and in cellular systems, its function in tissue biology, especially in the intestinal epithelium where it is expressed, remains unexplored. This work investigates this question in the context of gut epithelia development and morphogenesis.

Here we show that Myo1b is one of UNC-45A interactors, suggesting a role for myosin1b in the pathogenesis of UNC45A deficiency. Myo1b localizes at the apical brush border of intestinal epithelial cells in humans and loss of Myo1b in enterocyte like Caco-2 cells impairs epithelial morphogenesis. In zebrafish, genetic inactivation of Myo1b affects intestinal bulb fold formation revealing its conserved function during normal intestinal epithelia development.

## Results

### Myo1b is expressed in the gut epithelium and concentrates apically

Myo1b gene expression and protein localization were analyzed in intestinal epithelial cells. Myo1b was detected by Western Blot and immunofluorescence in the human epithelial colorectal Caco-2 cells (Fig. 1A-B). It accumulated apically in polarised Caco-2 cysts, as demonstrated by its colocalisation with the F-actin marker phalloidin demonstarting a localisation in actin-rich area, microvilli and/or the subjacent terminal web (Fig. 1B). As this model is adenocarcinoma cells, this expression and localization patterns could be the result of the tumoral state. To investigate Myo1b distribution *in vivo*, we looked for the homologue of *myo1b* in the zebrafish *Danio rerio*. There is one single *myo1b* gene with several splicing isoforms in the current zebrafish genome assembly. The corresponding Myo1b protein shares 80% identity with the *Homo sapiens* and *Mus musculus* homologues (Fig. 1C). In order to determine the expression pattern of *myo1b* during zebrafish development, we performed whole mount *in situ* hybridization labelling with specific antisense probes. *Myo1b* transcripts were unambiguously detected at 3dpf in the digestive tract of zebrafish (Fig. 1D) coinciding with the onset of gut morphogenesis (Ng et al., 2005, Wallace et al., 2005). *Myo1b* transcripts were also observed at 5dpf (Fig. 1D) when the intestine becomes functional and compartmentalised in bulb, mid and posterior intestines. At this stage, the transcripts were restricted to the intestinal bulb, the anterior part of the gut that forms large folds. Due to the lack of a zebrafish specific antibody, endogenous Myo1b sub-cellular localisation could not be assessed in zebrafish larvae. However, expressing eGFP-tagged Myo1b revealed apical accumulation of the protein, as previously seen in Caco-2 cysts by immunofluorescence (Fig. 1E).

**Figure 1.**
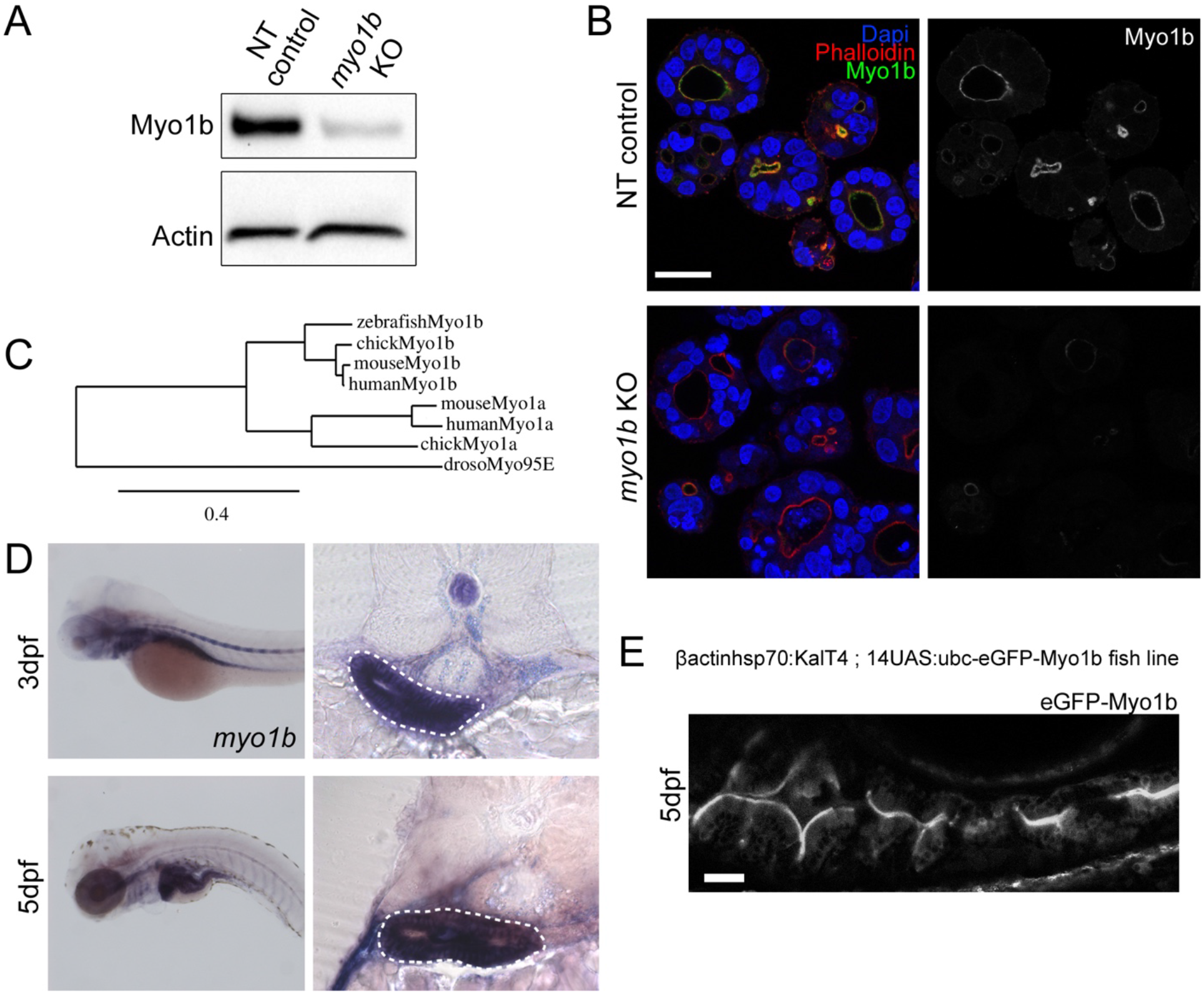
Myo1b expression and apical localisation in gut epithelial cells. **A-** Western blot analysis of Myo1b expression in extracts from non-targeted (NT) control and *myo1b* targeted Caco-2 cells (KO) using CRISPR/Cas9. **B-** Confocal sections of Caco-2 3D cultures stained for Myo1b, F-actin (phalloidin) and nuclei (Dapi). **C-** Phylogenetic tree based on protein sequence of zebrafish, chick, human and mouse Myo1b and Myo1a and drosophila Myosin95E. **D-** In situ hybridization for *myo1b* transcripts on 3 and 5dpf zebrafish larvae whole mounts (left panel) and cross-sections at the level of the intestinal bulb (right panels). On sections, the forming intestinal bulb is circled with white dashed lines. **E-** Live, longitudinal (antero-posterior axis) confocal section of the intestinal bulb of a 5dpf zebrafish larva expressing the transcription activator KalT4 driving the expression of the eGFP-Myo1b transgene under the control of an upstream activating sequence (UAS). The precise construction of the transgenes is annotated in the panel. Scale bars 30µm.

Myo1b is thus expressed in human intestinal epithelial cells and in the developing zebrafish intestinal bulb epithelium, and it preferentially localises apically in the brush border.

### *Myo1b* loss of function Caco-2 cysts show normal apico-basal polarization but altered luminal development

To address the function of Myo1b in enterocyte polarisation, *myo1b* was knocked-out using CRISPR/Cas9 in Caco-2 cells (*myo1b* KO, Fig. 1 A-B). The global apico-basal polarity of Caco-2 cysts was not affected in *myo1b* KO cells compared to controls as shown by the correct apical concentration of F-actin, pERM and villin (Fig. 2 A-B).

**Figure 2.**
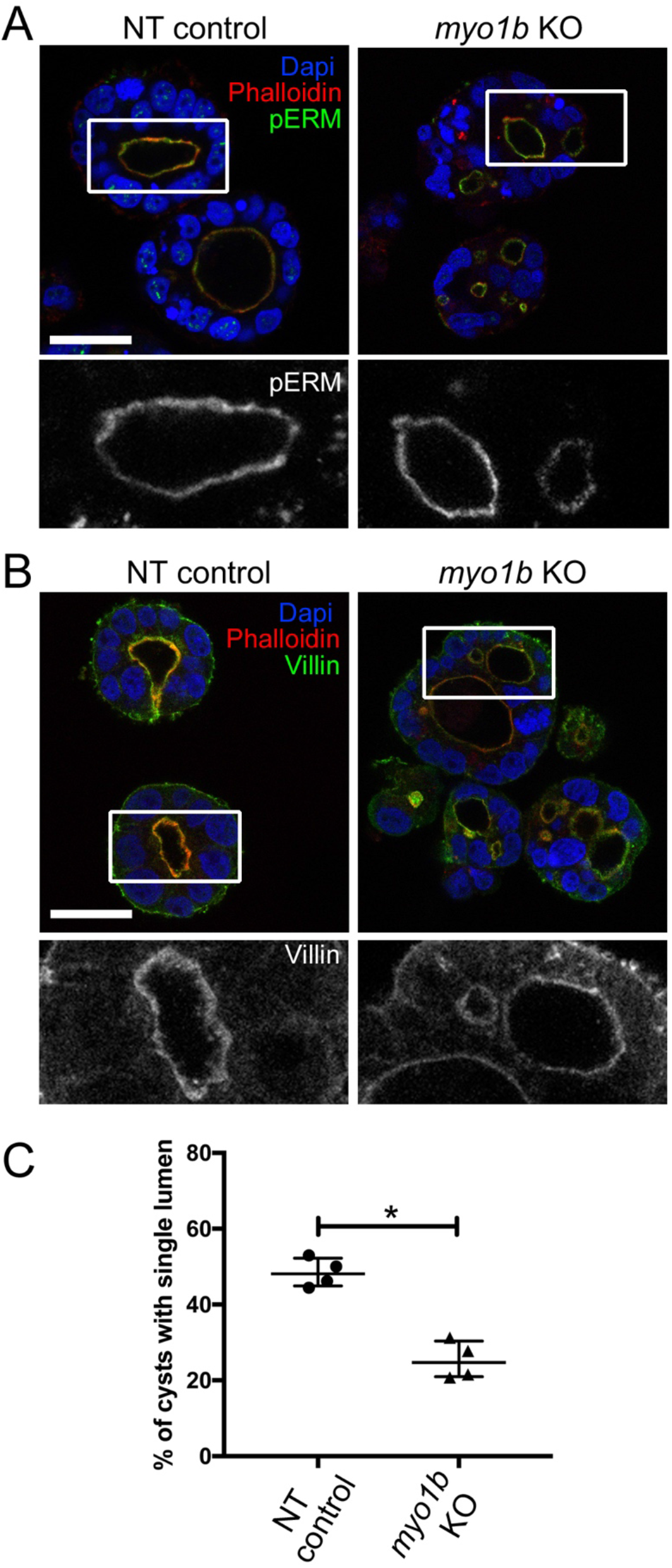
Enterocyte 3D cyst organization is affected in the absence of Myo1b despite normal apico-basal polarization. Confocal sections of NT control and *myo1b* KO Caco-2 3D cultures stained for the apical and microvilli markers phospho-Ezrin (pERM, **A**) and Villin (**B**). F-actin (phalloidin) and nuclei (Dapi) are stained, scale bars 30µm, boxed areas showed in insets are enlarged 2.5x. **C-** Quantification of the percentage of well-formed cysts with a single central lumen in NT control and *myo1b* KO Caco-2 3D cultures. Data represented are median and interquartile range from n=4 replicates, Wilcoxon test, *p<0.05.

Despite the absence of major polarisation defects, *myo1b* KO Caco-2 cells were more prone than controls to the formation of cysts with multiple lumen (Fig. 2). Indeed, *myo1b* KO cells showed a 50% drop in the percentage of well-formed cysts with single central lumen compared to controls (Fig. 2C).

### *Myo1b* loss of function has no major impact on epithelial cell differentiation of zebrafish intestinal bulb

To investigate the implication of Myo1b in intestinal epithelium morphogenesis *in vivo*, we turned to zebrafish as a good model for gut development. The zebrafish intestinal epithelium differentiates from 3 days post fertilisation (dpf) where it is essentially a flat monolayered tube. At 5dpf, epithelial folds are present, especially in the anterior most part of the gut, the intestinal bulb (Wallace et al., 2005). These folds are equivalent to the mammalian villi, and although no crypts are present in zebrafish, the region between folds will have a crypt-like role (Crosnier et al., 2005). First, we designed a splice blocking Morpholino (Myo1b-MO) that is efficiently preventing proper splicing of *myo1b* as determined by RT-PCR (supplemental Fig. 1A-B). At the concentration used, Myo1b MO displayed no overt phenotype, despite occasionally a slight heart oedema (supplemental Fig. 1C). To extend these results with a genetic loss of function model, we also generated a mutated allele at the *myo1b* locus using the CRISPR/Cas9 system, resulting in the insertion of a single base at the beginning of the open reading frame, as confirmed by sequencing (supplemental Fig. 1D). This leads to a premature stop codon and to the lack of detection of the protein by Western blot in gut lysates from adult homozygote mutants (*myo1b*-/-, supplemental Fig. 1E). As *myo1b* mRNA is maternally provided (supplemental Fig.1F), maternal contribution was suppressed by crossing *myo1b*-/-mothers. As for the MO injections, the resulting maternal-zygotic homozygous mutant larvae displayed no overt phenotype (supplemental Fig. 1C). In cross-sections (Fig. 3A), the intestinal bulbs of Myo1b MO and *myo1b*-/-larvae appeared smaller compared to controls. A significant reduction of the number of cells per cross-section was observed for both Myo1b MO and *myo1b*-/-intestinal bulbs at 3 and 5dpf compared to their respective controls (Fig. 3B). A reduction in the total cell number in the intestinal bulb could be the consequence of increased apoptosis or reduced cell proliferation. No significant difference with controls in the proportion of proliferative cells could be detected at 3 and 5dpf (supplemental Fig. 2A). A slight increase in the proportion of apoptotic cells could be detected at 5dpf but not at 3dpf (supplemental Fig. 2B). This later increase in apoptosis can however not account for the reduced cell number per section reported from 3dpf on and could more be a readout of increased cellular stress level upon prolonged absence of Myo1b, as reported after the KO of other Myosins 1 in mouse and drosophila (Hegan et al., 2007, Tyska et al., 2005). Using specific markers for secreting and absorptive cell lineages, defects in enterocyte differentiation could also be excluded (supplemental Fig. 2 C-D). Finally, the microvilli marker Villin appeared properly localised apically, lining the lumen together with F-actin (Fig. 3C), suggesting that apical polarity is not affected in Myo1b MO and *myo1b* -/- intestinal epithelium *in vivo*, as already shown in 3D Caco-2 cell cultures (Fig. 2B).

**Figure 3.**
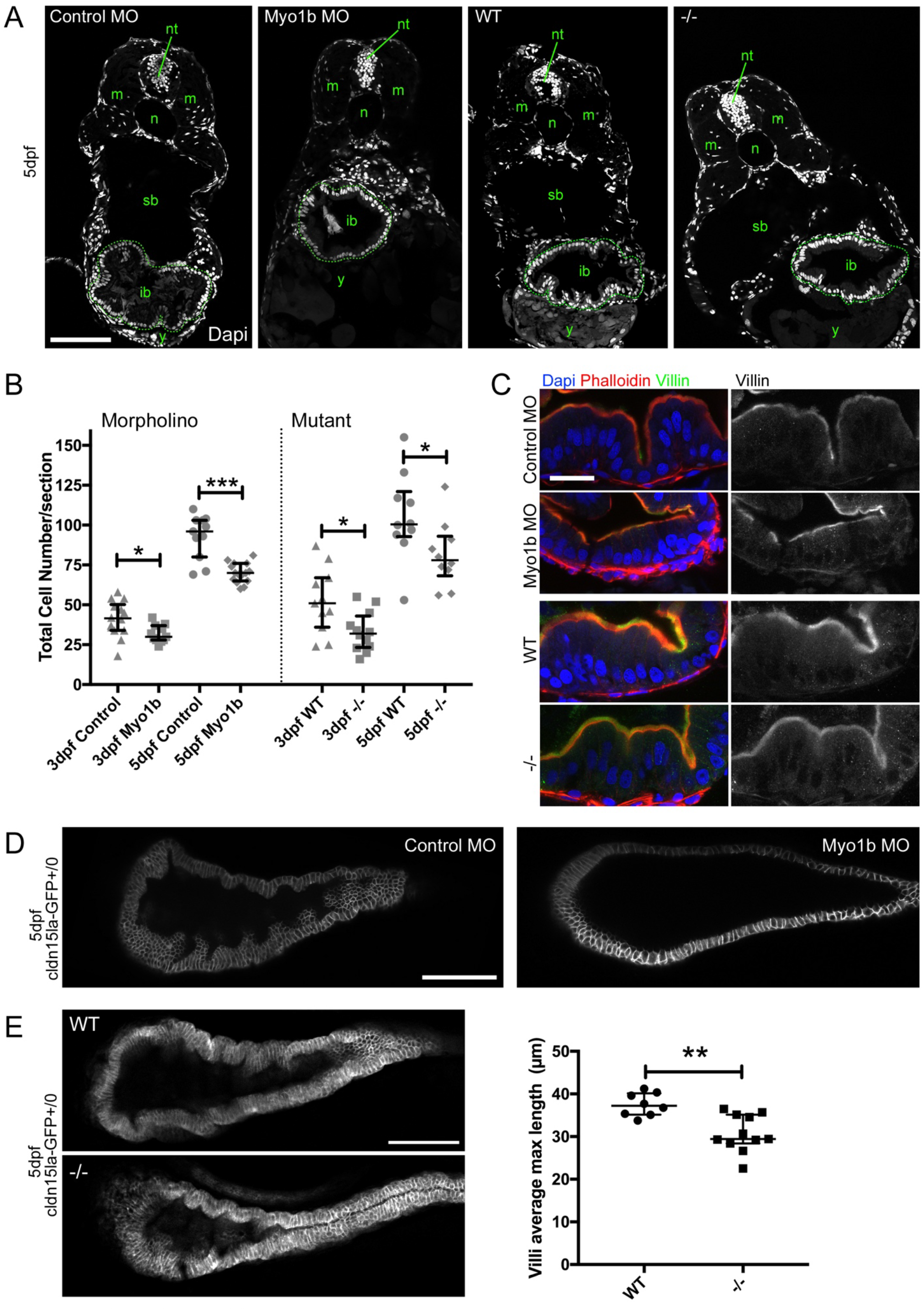
Myo1b knock-down and knock-out impair intestinal bulb fold development. **A-** Confocal single optical sections stained with nuclear labelling (Dapi) of 5dpf larvae injected with control and Myo1b Moprholinos (MO), and of 5dpf wild-type (WT) and *myo1b*-/-(-/-) larvae. ib intestinal bulb (circled with dashed lines), m muscles, n notochord, nt neural tube, sb swim bladder, y yolk. Scale bar=100µm. **B-** Quantifications from Dapi stained sections of the total number of cells per section at 3 and 5dpf in the four conditions. Data represented are median and interquartile range, n=10 to 15, Wilcoxon test, *p<0.05, ***p<0.001. **C-** Confocal optical sections of the intestinal bulb of 5dpf larvae in the four conditions stained for the microvilli marker Villin, F-actin (phalloidin) and nuclei (Dapi) showing the preserved apico-basal polarity of enterocytes when Myo1b is affected. Scale bar 20µm. **D-** Single confocal planes of live 5dpf larvae expressing Cldn15la-GFP injected with control MO (left) and Myo1b MO (right). Note the flat epithelium in the Myo1b MO condition. Scale bar=100µm. **E-** Single confocal planes of live 5dpf WT and *myo1b* -/- larvae expressing Cldn15la-GFP and quantification of the average length of the 3 longest folds per intestinal bulb analysed. Scale bar=100µm. Data presented are median and interquartile range, n_WT_=8 and n_-/-_=11, Wilcoxon test, **p<0.01.

### *Myo1b* loss of function zebrafish display MVID-like features in the intestinal bulb epithelium

To analyse in 3D intestinal bulb epithelium morphogenesis in zebrafish, we used the BAC line *cldn15la:cldn15la-GFP* that specifically labels the gut epithelium (Alvers et al., 2014). Both MO and KO intestinal bulbs revealed single continuous lumen suggesting that early steps of lumen fusion events were not affected (Alvers et al., 2014, Horne-Badovinac et al., 2001). However, in Myo1b MO larvae, the intestinal bulb epithelium appeared most of the time flat at 5dpf, not developing the expected folds observed in controls (Fig. 3D). Consistently with this phenotype, in the KO model, we detected a significant reduction in fold length in KO versus control samples (Fig. 3E). In an attempt to understand this milder phenotype in the mutant compared to the MO condition, we analysed potential compensation mechanisms by other myosins 1 performing RT-QPCR. On the 4 myosins 1 tested (*myo1ca, myo1cb, myo1d* and *myo1eb*), *myo1eb* showed a reproducible increase of on average 60% of the WT expression level in the mutant using EF1a (supplemental Fig. 1G) and RPL13a (not shown) as reference genes. *Myo1eb* is broadly expressed in early developmental stages but has restricted expression patterns after 2dpf (mostly branchial arches and pharynx) (Thisse and Thisse, 2004). A partial compensation of the loss of *myo1b* by an upreulation of *myo1eb* could thus potentially explain the subtler and more restricted phenotype of the mutant larvae compared to the knock-downs.

To further characterize the architecture of Myo1b-deficient intestinal bulb epithelium, a histological analysis by transmission electron microscopy (TEM) was performed on 5dpf larvae. It confirmed the affected folding of the intestinal bulb epithelium in MO and KO samples, and the preserved apico-basal polarity of enterocytes (Fig. 4A-B). Quantifying microvilli length and density did not reveal any significant defect resulting from Myo1b downregulation or absence, although packing looked less regular (Fig. 4C). In contrast, a darker sub-apical band was visible in Myo1b affected samples compared to controls, likely corresponding to modifications of the terminal web (Fig. 4C). Moreover, this ultrastructural analysis showed an important accumulation of vesicles in MO and KO samples compared to controls (Fig. 4B insets) suggesting defects in membrane trafficking. These TEM observations (epithelium folding impaired, modifications of the apical pole ultrastructure and trafficking defects) overall indicate that myo1b-deficient enterocytes display some microvillus inclusion disease-like features.

**Figure 4.**
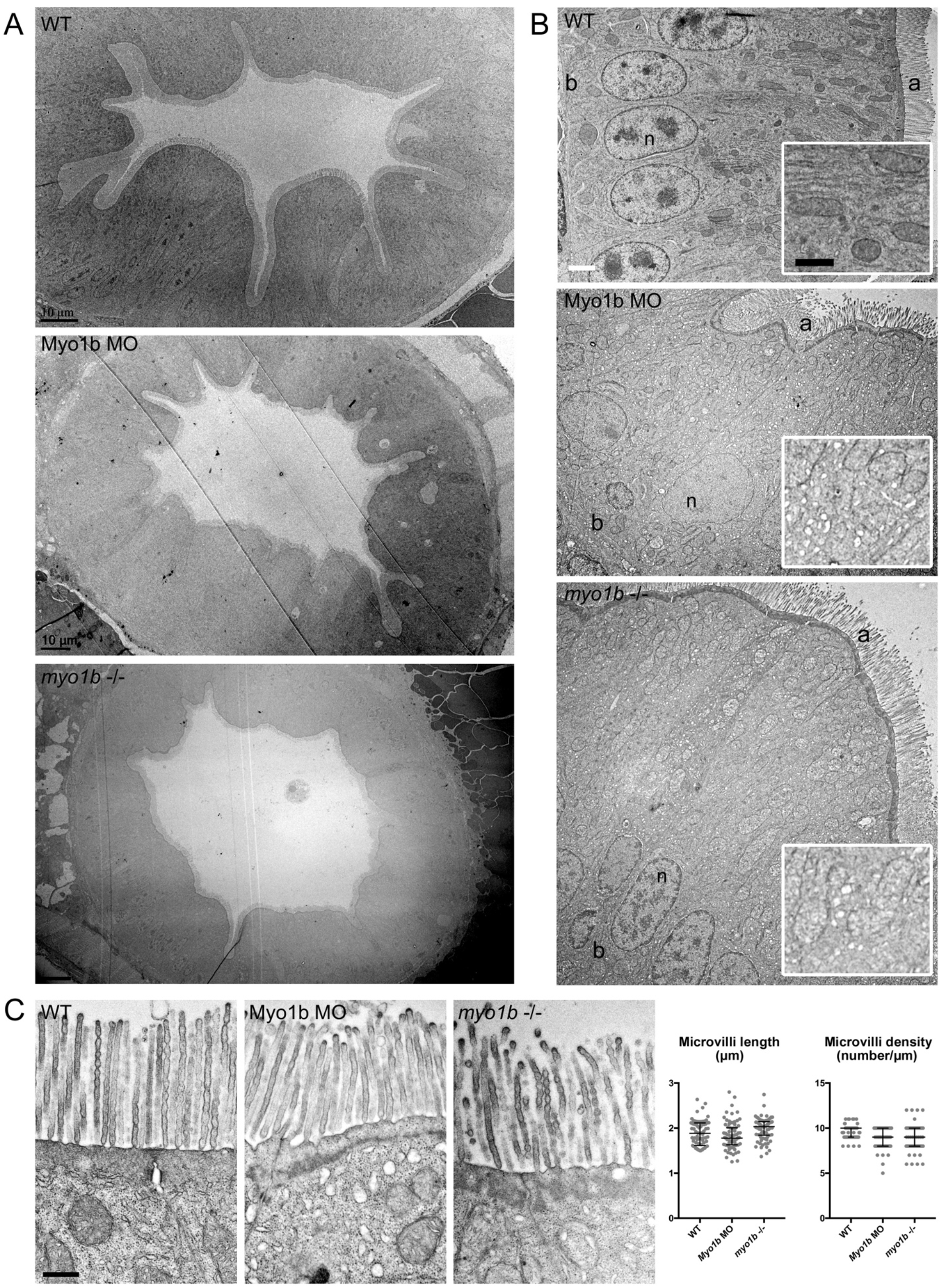
Electron Microscopy confirms folding defects and shows affected trafficking. Transmission electron micrographs of sections of intestinal bulbs from WT, Myo1b MO and *myo1b-/-* 5dpf larvae presenting a general view of the folds of the epithelium (**A**, scale bars 10µm) and of the apico-basally polarized enterocytes (**B**, scale bar 2µm; b basal, a apical, n nuclei). Insets in **B**, show higher magnifications of the cytoplasm region to highlight the accumulation of vesicles in Myo1b MO and *myo1b*-/-samples, scale bar 1µm. **C-** Transmission electron micrographs of sections of intestinal bulbs from WT, Myo1b MO and *myo1b-/-* 5dpf larvae illustrating the organization of the brush border, and quantifications of the average length and density of the intestinal microvilli in the different conditions. Data presented are median and interquartile range, n_length_=77, n_density_=75 per condition. Scale bar 500 nm.

### Myo1b is destabilized when Unc45A is affected

Loss of function (LOF) mutations in the chaperone *UNC45 homolog A* have recently been associated with rare human genetic syndromes notably presenting intestinal disorders, including chronic diarrhea and villous atrophy of variable penetrance (Esteve et al., 2018). A pull-down assay performed to detect potential Myo1b partners identified UNC45A as the most abundant protein interacting with Myo1b in a mouse neuronal cell model (Supplementary table 1). This interaction was confirmed by the reverse experiment using UNC45A as bait in the colorectal Caco-2 cells (Duclaux-Loras *et al*., submitted).

Considering this interaction, we investigated the impact of *UNC45A* depletion on Myo1b expression. *UNC45A* depleted Caco-2 cells showed reduced Myo1b levels by immunofluorescence (Fig. 5A). To detect aggregation-prone proteins normally sent to degradation, the proteasome machinery was blocked using the proteasome inhibitor MG132. MG132 induced the appearance of protein aggregates both in control and UNC45A KO Caco-2 cells (Fig. 5B). Myo1b staining partly co-localised with the aggresomes in the UNC45A KO condition (Fig. 5B). This result suggests that Myosin1b proper folding requires the chaperone UNC45A.

**Figure 5.**
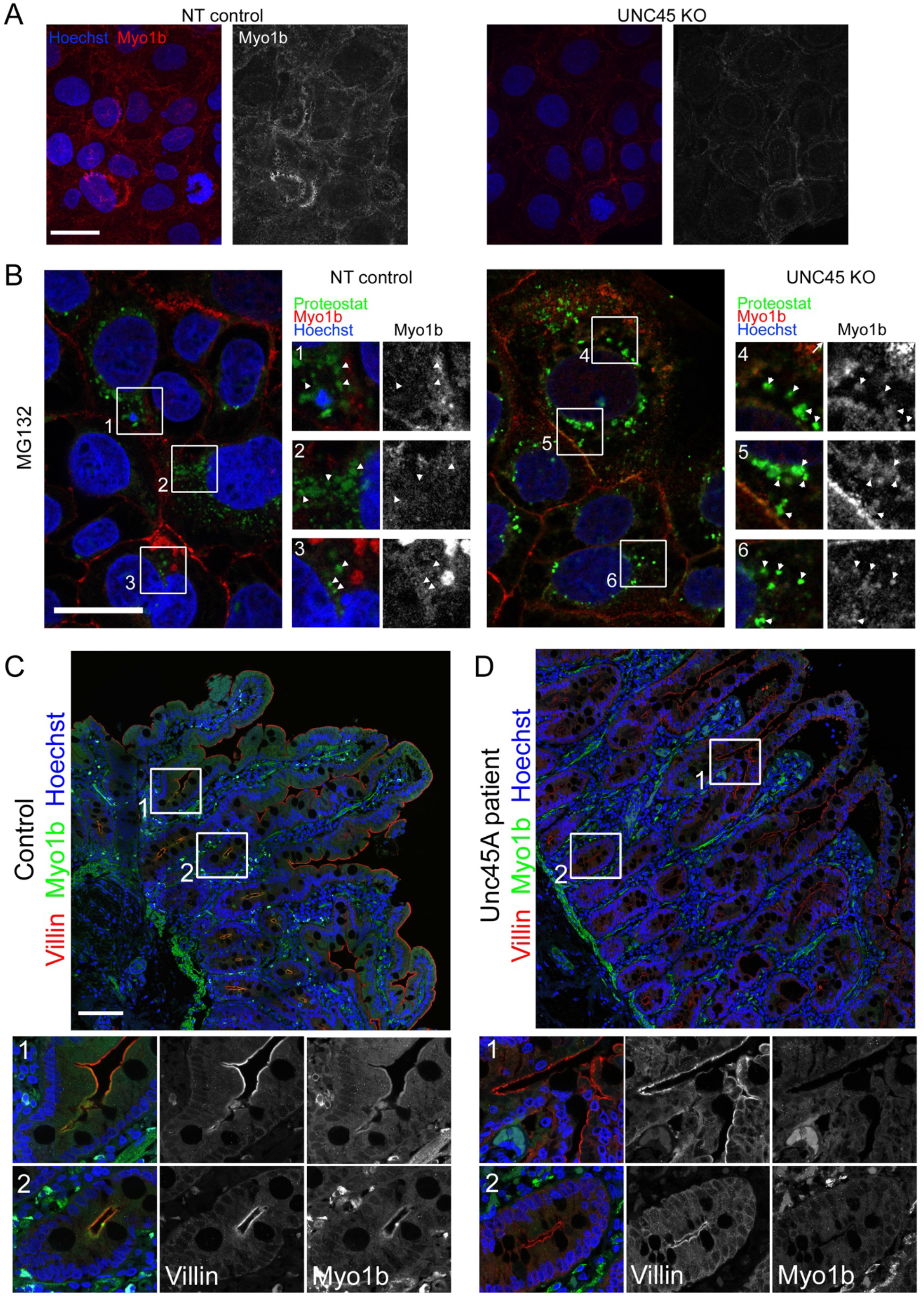
Myo1b expression is destabilized in UNC45A depleted cells and in biopsies from UNC45A mutated patients. **A-** Immunohistochemistry analyses of Myo1b in non-targeted (NT) control and UNC45A deficient (KO) Caco-2 cells show decreased Myo1b levels. Pictures are maximal projections of confocal stacks, Hoechst labels nuclei, scale bar 30µm. **B-** Confocal sections of NT control and UNC45A KO Caco-2 cells treated with the proteasome inhibitor MG132 and stained for Myo1b and the aggresome probe Proteostat. Hoechst labels nuclei, scale bar 30µm. Boxed areas showed in insets are enlarged 2x, arrowheads point at Proteostat-labelled protein aggregates and highlight colocalisation with Myo1b proteins in UNC45A KO cells. **C**,**D-** Confocal sections of a human biopsy from a healthy patient (control, C) and from a UNC45A LOF patient (D) immuno-labelled for the microvilli marker Villin and for Myo1b; Hoechst labels nuclei, scale bar 100µm. Boxed areas showed in insets are enlarged 3x, and highlight the apical localisation of Myo1b in control tissue (C) at the base of the villi (1) and in crypts (2), which is essentially lost in the UNC45A LOF tissue (D).

We finally looked at Myo1b expression in duodenal biopsies from control and *UNC45A* loss of function patients. In control biopsies, the microvilli marker Villin was expressed apically all along the epithelium and Myo1b was detected apically at the base of the villi and in crypts, partially colocalising with Villin (Fig. 5C). In *UNC45A* patients, Villin was still localised apically whereas Myo1b was barely detectable (Fig. 5D). In conclusion, Myo1b protein level is decreased in *UNC45A*-depleted cells and in duodenal biopsies from an *UNC45A*-mutated patient.

## Discussion

This work identifies Myo1b, an actin motor, as an unexpected player in the regulation of the morphogenesis of the intestinal epithelium during gut development. In zebrafish, we report defects in epithelial folding and villous atrophy when Myo1b is impaired. This phenotype is similar to the ones reported in zebrafish and in human patients with loss of function mutations in *UNC45A*. We show that Myo1b interacts with UNC45A and that Myo1b is destabilised in absence of UNC45A.

For this study, we analysed both *myo1b* mutant- and Morpholino-induced phenotypes. It is now well established that Morpholino knock-downs often result in more severe overt phenotypes than the corresponding knock-outs, at least partially due to the induction of genetic compensation mechanisms in the mutants (Kok et al., 2015, Rossi et al., 2015). In the *myo1b* null case, we observed a more subtle outcome than the Mopholino, which could be due to compensation mechanisms (El-Brolosy and Stainier, 2017) as supported by the RT-QPCR of *myo1eb*. Here, the intestinal bulb phenotypes observed with both approaches converged on reduced cell number of intestinal bulb sections and impaired epithelial folding.

Up to now, myosins 1a, c, d and e had been identified in intestinal brush borders constituting the apical pole of differentiated enterocytes (Benesh et al., 2010). Several class I myosins have been implicated in the maintenance of intestinal epithelial differentiated state. Myo1a, which is associated with the highly organised actin network of differentiated enterocytes in mammals (Revenu et al., 2012, Tyska et al., 2005), but seems to lack in the zebrafish and *Drosophila* genomes, is important for enterocyte polarity and participates in the structure and composition of the brush border (Mazzolini et al., 2012, Tyska et al., 2005). The phenotype of the *myo1a* KO mice is however mild, with reports of clear compensations by other class I myosins (Benesh et al., 2010, Tyska et al., 2005). Likewise, two of the known class I myosins in *Drosophila* are also localised in the apical pole of differentiated enterocytes and *Drosophila* Myo61F is necessary for the stability of enterocyte apical organisation (Hegan et al., 2007). We report the expression of another myosin 1 in the gut epithelium, Myo1b, and its apical localisation in enterocytes of Human biopsies, of zebrafish intestinal bulb and in Caco-2 cells. Myo1b localisation in microvilli had previously been reported in kidney epithelial cells (Komaba and Coluccio, 2015). In Human biopsies, Myo1b is expressed at the base of the villi and in crypts suggesting a specific role in the proliferative compartment and not in enterocyte differentiated state. We did not detect global polarisation defects at the cell level or impaired differentiation in absence of Myo1b, whereas both the Caco-2 3D model and the zebrafish model demonstrate morphogenetic defects at the tissue level, respectively single lumen formation and folding.

Proliferation, apoptosis or differentiation defects do not account for the reduced cell number observed on transverse sections of the intestinal bulb from 3dpf. As the sections give a 2D overview of a 3D organ, this reduced cell number is thus likely the readout of the different organisation in space of the epithelium. A specificity of the zebrafish intestinal bulb is the early folding of the epithelium that remains pronounced to adulthood (Ng et al., 2005). The reduced folding and villi formation in the mutants and morphants are clear signs of a different architecture of the tissue. Also the mechanisms underlying intestinal epithelium folding are not yet fully understood, the impact of tension and forces at the cell and tissue level driving compression, cell intercalation and invagination through apical constriction have been investigated in other tissues during development (Mammoto and Ingber, 2010). Myosins are central in the control of actin cytoskeleton dynamics and in force generation (Reymann et al., 2012). In particular, Myo1b deforms membranes and participates in organelle formation and trafficking (Almeida et al., 2011, Coudrier and Almeida, 2011). It also remodels the actin cytoskeleton (Iuliano et al., 2018, Pernier et al., 2019, Prosperi et al., 2015). Its roles in membrane traffic and in the dynamic organisation of actin structures make it a plausible actor in the morphogenesis of the gut epithelium. The electron microscopy data show a strong accumulation of intra-cellular vesicles in Myo1b mutant and Morpholino tissues suggesting impaired trafficking, in agreement with its role in the formation of post Golgi carriers and protein transport at the level of multivesicular endosomes (Almeida et al., 2011, Salas-Cortes et al., 2005). Electron microscopy also reveals modifications of the terminal web area, the apical actin belt linking adherens junctions in the epithelium, in agreement with its role in actin dynamics. Myosin 1b function on actin dynamics and consequently on membrane remodelling and membrane trafficking must impact cell and tissue mechanics (Buske et al., 2012), and this way contributes to impaired intestinal epithelial folding in the absence or down-regulation of *myo1b*. Myo1b restricted localisation at the base of the villi in human biopsies may indicate a specific mechanical role in crypts morphogenesis.

Villous atrophy is a phenotype associated with various intestinal disorders including some rare hereditary syndroms presenting congenital diarrhea like microvillous inclusion disease (MVID). Mutations in MyosinVb are the main cause of MVID (Muller et al., 2008) and have notably been associated with defective trafficking. A zebrafish mutant of *myoVb* develops a flat intestinal epithelium (Sidhaye et al., 2016). Recently, loss of function (LOF) mutations in the chaperone *UNC45A* were reported in families presenting complex phenotypes including congenital diarrhea and several degrees of villous atrophy (Esteve et al., 2018). A zebrafish mutant for *UNC45A* also exhibit loss of intestinal epithelium folding (Esteve et al., 2018). UNC45A is a chaperone participating in the conformational maturation of, among others, some Myosins (Barral et al., 2002, Lee et al., 2014, Lehtimaki et al., 2017). Our results in human cell lines and in patient samples demonstrate a strong reduction in the protein level of Myo1b in absence of a functional UNC45A variant, probably due to the degradation of misfolded and destabilised Myo1b. Myo1b proper conformational folding would thus require the chaperone UNC45A. The intestinal phenotypes associated with LOF mutations in *UNC45A* could hence partly be the consequence of the reduced protein level of Myo1b. In conclusion, Myo1b contributes to gut morphogenesis and appears as a potential player in the complex intestinal phenotype of the UNC45A LOF syndrome.

## Materials and Methods

### CRISPR- Cas9 genome editing of MYO1B in Caco2 cells and 3D culture

The lentiCRISPRv2 plasmid was a gift from F. Zhang (Massachusetts Institute of Technology, Boston, MA; plasmid no. 98290, Addgene). The single-guide RNAs (sgRNAs) were designed using the CRISPR Design Tool (Massachusetts Institute of Technology) and cloned into the BsmbI site. sgRNA sequences: for-5-CACCGATCCCTACGAGATCAAGATA −3, rev-5-AAAC TA TCT TGA TCT CGT AGG GAT C −3. Following production of lentiviral particles, the lentiCRISPRv2 plasmids were transduced in Caco2 cells. Positively transduced cells were selected by puromycin (10μg/ml). For 3D culture, CaCo2 cells were resuspended at a concentration of 10^4^ cells/mL in DMEM (Gibco) with 20% FCS containing 4% Matrigel (BD Biosciences) and 2.5 10^4^ cells/well were plated in 8-wells chamber slide IBIDI (Biovalley), previously precoated with 100 µL of Matrigel. Cells were grown for 5 days to obtain cysts. To detect aggregation-prone proteins, the proteasome inhibitor MG132 (Sigma-Aldrich) was added overnight in the culture medium at a concentration of 10 μM.

### Western blot

Caco-2 cells were lysed in RIPA buffer (Sigma) supplemented with 1X proteinase inhibitor cocktail mix (Roche, Sigma). Adult zebrafish guts were dissected on ice and mechanically lysed in 200μL lysis buffer (10 mM HEPES + 300 mM KCl + 5 mM MgCl_2_ + 0,45% triton X100 + 0,05% Tween20, pH7) with 10 mM ATP and Complete protease inhibitor (Roche). 40 μg of extracts in Laemmli buffer were loaded on a 4-12% polyacrylamide gradient concentration gel (ThermoFisher). Primary antibodies used were mouse anti-tubulin (1:12000, Sigma), rabbit anti-ratMyo1b (1,8 μg/μL, 1:500, (Salas-Cortes et al., 2005), anti GAPDH (1:1000, Cell Signaling).

### Phylogenetic analysis

The Myo1b and Myo1a homologues in *Danio rerio, Homo sapiens, Mus Musculus, Gallus gallus and Drosophila melanogaster* were obtained from NCBI HomoloGene. Protein sequences were aligned and a phylogenetic tree was assembled using the online ‘One Click’ mode at Phylogeny.fr (Dereeper et al., 2008).

### Molecular Cloning

The *βactinhsp70:KalT4;cmlc2:eGFP* construct was generated by combining four plasmids using the Multisite Gateway system (Invitrogen): p5E-bactinhsp70, pME-KalT4, p3E-polyA and pDEST-cmcl2:eGFP containing Tol2 sites (Kwan et al., 2007). The βactin promoter was cloned into the pCR-bluntII-TOPO vector (Invitrogen) and then inserted in the p5E-MCS using KpnI and XhoI restriction sites. The 3’ 638bp of the zebrafish hsp70 promoter (Dalgin et al., 2011) was inserted into this p5E-βactin vector linearized with XhoI using the Gibson Assembly Cloning Kit (New England Biolabs). The optimised Gal4 KalT4 (Distel et al., 2009) was amplified and inserted in a pDONR221 using the Multisite Gateway system (Invitrogen). To generate the *14UAS:ubc-eGFP-Myo1b vector*, eGFP-Rat Myo1b cDNA was amplified from a previously published plasmid (Prosperi et al., 2015). It was inserted into the pT1UciMP plasmid containing 14xUAS-ubc and Tol1 sites (Horstick et al., 2015) linearized with Xho1 using the Gibson Assembly Cloning Kit (New England Biolabs).

### Zebrafish (*Danio rerio*) husbandry

Wild-type Tupfel long fin zebrafish strains were used and raised according to standard protocols. Stable transgenic lines were generated by injection of the plasmids with *tol2* or *tol1 transposase* mRNA at 25ng/µL in one-cell stage zebrafish embryos. The transgenic BAC line claudin15-like-a fused to GFP (cldn15la:cldn15la-GFP) was kindly provided by Michel Bagnat (Alvers et al., 2014).

For live-imaging, larvae were anaesthetised in 0.02% MS-222 and immobilised in 1% low melting point agarose. Imaging was performed on a Zeiss LSM 780 confocal. All animal procedures were performed in accordance with French and European Union animal welfare guidelines.

### *In situ* hybridization

*In situ* hybridizations (ISH) were performed on larvae treated with 1-phenyl-2-thiourea (PTU, Sigma-Aldrich) and fixed in freshly made 4% paraformaldehyde (PFA) 2-4h at RT and stored in 100% methanol at −20°C. After rehydration, larvae were treated with proteinase K (20 µg/ml; Roche diagnostics) at RT for 1h (3dpf) or 2h (5dpf) and fixed again in 4% PFA at RT for 20min. Digoxigenin-labelled antisense and sense RNA probes were synthesized by *in vitro* transcription using DIG-labelled UTP according to the manufacturer’s instructions (DIG RNA labelling kit, Roche). Primers used were as follow: *myo1b* sense: CAA TAT GAT AGG GGT AGG GGA CAT G; antisense: TGG TTT GAA CTC AAT ATT TCC CAG C. Anti-DIG antibody conjugated to alkaline phosphatase allowed detection of hybridized riboprobes according to the manufacturer’s instructions (Roche).

### *Myo1b* zebrafish mutant generation with CRISPR/Cas9

The sgRNA sequence (sgB: CTTCTGACAAGGGCTCTAGG) was cloned into the BsaI digested pDR274 vector (Addgene). The sgRNAs were synthesized by *in vitro* transcription (Megascript T7 transcription kit, Ambion). After transcription, sgRNAs were purified using the RNAeasy Mini Kit (Quiagen). For injections at one cell stage, the synthesized sgB was injected at 300ng/µL after 5-minute incubation at RT with Cas9 protein (NEB) at 2µM final in 20mM Hepes-NaOH pH 7.5, 150mM KCl (Albadri et al., 2017). Injected embryos were grown to adulthood and crossed with wild-types to identify founders. Pools of 20 embryos per clutch were lysed in NaOH 50mM at 95°C for at least 30min. PCR was performed on lysates to amplify the genomic region targeted by the sgB with primers forward 5’GGGTGTTGTTCAGCGATGGA and reverse 5’ATAGATCTCATTGTGATCGA using Phusion High-Fidelity DNA polymerase (Thermo Scientific). The amplicons were cloned in pCR-bluntII-TOPO vector (Zero Blunt Topo PCR cloning kit, Invitrogen) and sequenced (GATC Biotech) to identify indels and the corresponding founder fish. Sequences were analysed using Geneious. After selection of the founder, genotyping of the line was performed by PCR on fin clips with primers 5’AGATGAATGCAAGCAAGCCATT and 5’ATACGATCTGATTGTGATCGAATCGCT. The resulting product was digested with restriction enzyme FspBI, the site of which is lost in the mutant, resulting in 2 fragments (208 and 66bp) for the WT allele and only one (275bp) for the mutated allele.

### Morpholino oligonucleotide design and injections

*Myo1b* splice blocking morpholino was designed to target the splice donor site downstream of exon 22 (Myo1b-MO, 5’-ATGAGAAACTGTGTTCATTACCTGG). Experiments were performed in parallel with a standard control-MO (5’-CCTCCTACCTCAGTTACAATTTATA) with no target in the zebrafish genome. Morpholinos (Gene Tools) diluted at 1mM in water, were injected in 1-cell stage embryos at a final concentration of 0.2mM. To validate Myo1b-MO knock-down, RT-PCR was performed on 3dpf larvae. Total RNA of 50 larvae was prepared with TRIzol and TURBO DNAse-free reagents (Invitrogen). mRNA (1μg) was retro-transcribed using oligo(dT) primers and the SuperScript III First-Strand Synthesis System (Invitrogen). To amplify the region targeted by the MO (supplemental Fig. 1A), PCR was performed on the cDNA with two different forward primers (primer 1 in exon21 was: 5’GGCTGCGATATTCTTGCCTCC, primer 2 at the edge of exon22 and the targeted intron was: 5’TCTTTCATTCGTGGATGGAAGGCC) and the reverse primer 5’AACCCAGGTAATGAACACAGTTTCTAT. PCR products were run on a gel (supplemental Fig. 2B) and bands were gel purified (Macherey-Nagel), inserted in a pCR-bluntII-Topo vector (Invitrogen) and sent for sequencing (GATC) to assess intron retention.

### Quantitative RT-PCR

For each experiment, total RNA was prepared from 3 pools of 50 embryos per phenotype with TRIzol reagent and the TURBO DNA-free kit (ThermoFisher Scientific). RNA (1μg) was retro-transcribed using random primers and the SuperScript III First-Strand Synthesis system (ThermoFisher Scientific). For RT-QPCR, the SYBR Green PCR Master Mix (ThermoFisher Scientific) was used according to the manufacturer’s instructions and PCR were performed on an ABI PRISM 7900HT instrument. Experimental triplicates of each sample were averaged and the relative expression level quantified with the log2ΔCT method using EF1a and RPL13a reference genes. Shown are values normalised on the wild-type samples.

### Immunohistochemistry

Caco-2 cells and cysts were fixed in 4% PFA 30min at 37°C and washed with PBS. They were permeabilised in PBS, 0.2%TX100, 1%BSA for 5min at RT and blocked in PBS 3%bBSA for 1-2 hours at RT. Primary antibodies used were rabbit anti-ratMyo1b (1,8 μg/μL, 1:100, (Salas-Cortes et al., 2005), mouse anti-Villin clone ID2C3 (1:300, (Robine et al., 1985), rabbit anti-pERM (1:100, abcam ab47293). After washes, they were incubated with Alexa Fluor 488 secondary antibody (Molecular probes), phalloidin-Alexa Fluor 568 and Dapi. To assess protein aggregation, the Proteostat Aggresome Detection Kit (ENZO, ENZ-51035) was used. Briefly, after the primary antibody rabbit anti-ratMyo1b, cells were incubated for 30min at RT with goat anti-rabbit-Alexa Fluor 635 antibody (1:400, Molecular probes), Proteostat 1:400 and Hoechst 1:800 (ENZO) in PBS 3%BSA.

Zebrafish larvae were fixed for 2h at room temperature in 4% PFA and incubated in 30% sucrose/0.1% PBST overnight at 4°C. They were then frozen in Tissue-Tek OCT (Sakura) at −80°C and sectioned using a Cryostat (Leica). Zebrafish larvae sections were incubated in blocking buffer (10% serum in PBST, PBS 0.1%Tween20) and with mouse anti-Villin clone ID2C3 (1:300), mouse 2F11 antibody (1:100, Abcam ab71286) or 4E8 (1:100, Abcam ab73643) overnight at 4°C. After washes with PBST, they were incubated with Alexa Fluor 488 secondary antibody (Molecular probes), phalloidin-Alexa Fluor 568 and Dapi. To assess apoptosis, TUNEL assay was performed with reaction solutions from ApopTag Red In situ Apoptosis detection kit (Millipore) according to the manufacturer recommendations. To assess proliferation, larvae were injected in the yolk with 10mM 5-ethynyl-2’-deoxyuridine (EDU) in 1% DMSO and incubated in 100µM EdU, 0.4%DMSO for 20 hours after injection. Animals were fixed at indicated time and processed according to the Click-iT EdU Imaging Kit (Invitrogen).

Paraffin embedded sections of intestinal tissues from a UNC45A deficient patient and biopsies from controls were obtained for diagnosis or therapeutic purposes. Duodenal biopsies were routinely fixed in 4% buffered formalin for 24 hours and paraffin-embedded. Sections were heated for 1hr at 65°C and paraffin was removed by two 5-min washes in xylene. Sections were then hydrated with ethanol solutions of decreasing concentrations. Unmasking of the epitopes was performed at 100°C for 20 min in Citrate-based Antigen Unmasking Solution (Vector Laboratories). Sections were incubated for 30 min at room temperature in blocking buffer (3% BSA in PBS) and then overnight at 4°C with anti Myo1b antibody (1:200, Novus Biologicals NBP1-87739) in blocking. After washes with PBST, sections were incubated with goat anti rabbit Alexa Fluor 488 antibody (Molecular probes), phalloidin-Alexa Fluor 568 and Dapi for 2hrs at RT.

After extensive washes and mounting in Vectashield (Vector Lab), all stainings were imaged on a LSM780 confocal microscope (Zeiss). Images were processed and numbers of cells quantified using ImageJ.

### TEM analysis on zebrafish larvae

5dpf larvae were collected and stored at 4°C in Trump’s fixative. Enhanced chemical fixation was performed in a mix of 4% PFA with 2.5% glutaraldehyde in 0.1 mol/L cacodylate buffer overnight at 4°C. A 1.5-hour incubation in 1% OsO4 was followed by a 1.5-hour incubation with 2% uranyl acetate at ambient temperature. Larvae were then dehydrated through graded ethanol solutions, cleared in acetone, infiltrated, and embedded in Epon-Araldite mix (EMS hard formula). We used adhesive frames (11560294 GENE-FRAME 65 µL; Thermo Fisher Scientific) for flat-embedding, as previously described (Kolotuev et al., 2012), to gain better anteroposterior orientation and sectioning. Ultrathin sections were cut on an ultramicrotome (UC7; Leica Microsystems) and collected on formvar-coated slot grids (FCF2010-CU, EMS). Each larva was sectioned transversally in five different places in intestinal bulb with ≥20 μm between each grid to examine the sample over a large region. Each grid contained at least 4-6 consecutive sections of 70 nm. TEM grids were observed using a JEM-1400 transmission electron microscope (JEOL) operated at 120 kV, equipped with a Gatan Orius SC1000 camera (Gatan) and piloted by the Digital Micrograph program. Microvilli length and density were quantified using Fiji on TEM pictures of at least 50 MV from 25 enterocytes of 3 larvae per condition.

### Pull Down assay

10^6^ N1E115 cells were transfected with pEGFP Myo1b (Salas-Cortes et al., 2005) and lysed in TRIS 150mM, Nacl 150mM, EDTA 1mM, EGTA 1mM, ATP 10 mM, 10% glycerol, 1mM DTT, 0,5% triton and protease inhibitor 24 hours after transfection. The lysate was then incubated with 15 ml of GFP trap Beads (Chromotek) overnight. After washing the beads were resuspended in water and treated for mass spectrometry analysis.

### Statistical analysis

The numbers of cells reported are coming from manual counting. No sample was excluded from the analysis, except for the total cell number per section where we made sure to analyse samples displaying single cell layers through the whole gut cross-sections and not the side of some villi. The sample size (n=) is defined as the number of larvae analysed (one section per larva). For statistical analysis, we applied the non-parametric Wilcoxon-Mann Whitney test.

## Acknowledgements

The authors thank the Del Bene team for fruitful suggestions and discussions and members of the Institut Curie zebrafish facility. The authors acknowledge all members from the PICT-IBiSA Lhomond Imaging Platform (UMR144) and the Cell and Tissue Imaging Platform of the Genetics and Developmental Biology Department (UMR3215/U934) of Institut Curie, member of France-Bioimaging (ANR-10-INSB-04), for help with light microscopy and the electron microscopy unit of the MRic facility (Rennes, France).

## Contributions

MP generated the KO cells. JS and CR generated the *Myo1b* null allele in zebrafish. MR generated the stable zebrafish transgenic lines. CR performed immunofluorescence stainings. KD, JV and MR performed ISH. MP, MR, CR and RDL performed WB. KD and CR did the RT-qPCR analysis. JS and JV genotyped the mutants. ON and GM did the TEM and analysis. MP, RDL and CL prepared cell culture samples. PL generated preliminary data with Morpholinos. MTP and EC performed the pull-down assay. CR analysed the results and wrote the paper. FDB, EC and NCB supervised the work. MR, MP, GM, EC and FDB edited the manuscript.

## Supplementary Figures and Table

**Figure S1.**
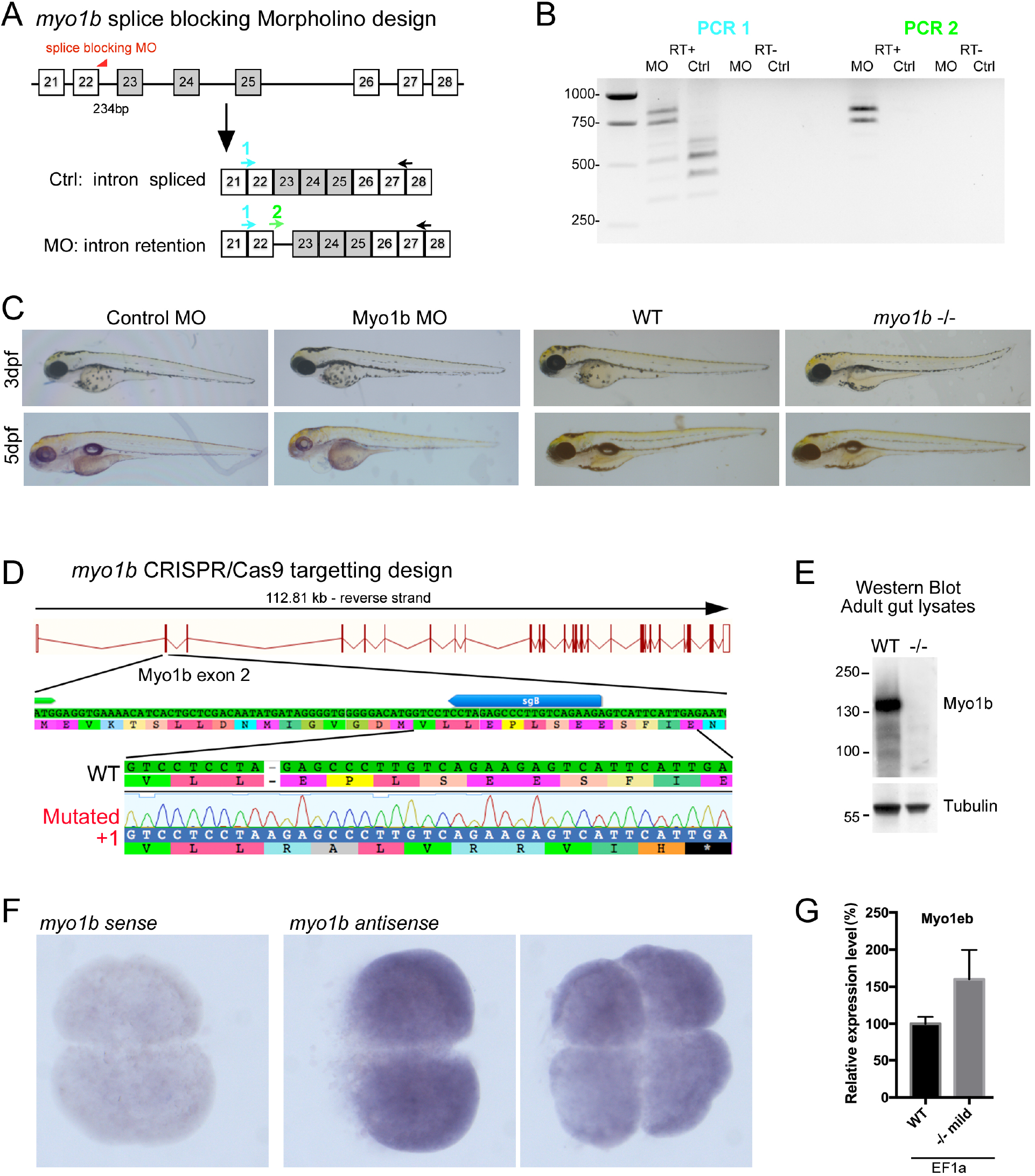
Myo1b Morpholino and CRISPR mutant design and validation. **A-** Schematics of the design and **B-** DNA gel of the RT-PCR performed to control Myo1b-MO knock-down efficiency. Higher bands amplified in PCR1 MO compared to control and bands amplified in PCR2 MO correspond to Myo1b cDNA retaining the intron targeted by Myo1b-sMO, as verified by sequencing. In B, the multiple bands amplified, both in control and MO conditions, correspond to expected splicing variants of exons 23, 24 and 25 (highlighted in grey) as checked by sequencing. RT-is the control RT without superscript compared to RT+. **C-** Bright field pictures of 3 and 5dpf larvae presenting the phenotypes of control and Myo1b Morpholinos, WT and *myo1b*-/-larvae. **D-** Schematics of CRISPR/Cas9-mediated gene disruption at the *myo1b* genomic locus. The sgRNA (sgB, blue arrow) was targeting exon 2 downstream the start codon (ATG, green arrow). Compared to the WT sequence, the mutated allele displayed an insertion of 1bp generating a frame shift from amino-acid 21 and a premature STOP codon after 29 amino-acids. **E-** Western Blot with antibodies against Myo1b and Tubulin on lysates of dissected guts from WT and -/- adults. **F-** Negative control with a sense probe and in situ hybridisation with a *myo1b* anti-sense probe on wild-type embryos at 2 and 4 cell-stages showing maternal contribution for *myo1b* mRNA. **G-** RT-QPCR of *myo1eb* expression at 3dpf with EF1a used as reference gene, normalised on expression of the WT samples. Shown are mean and sem (WT=100.0±9.2, -/- =160.1±39.5, n=6).

**Figure S2.**
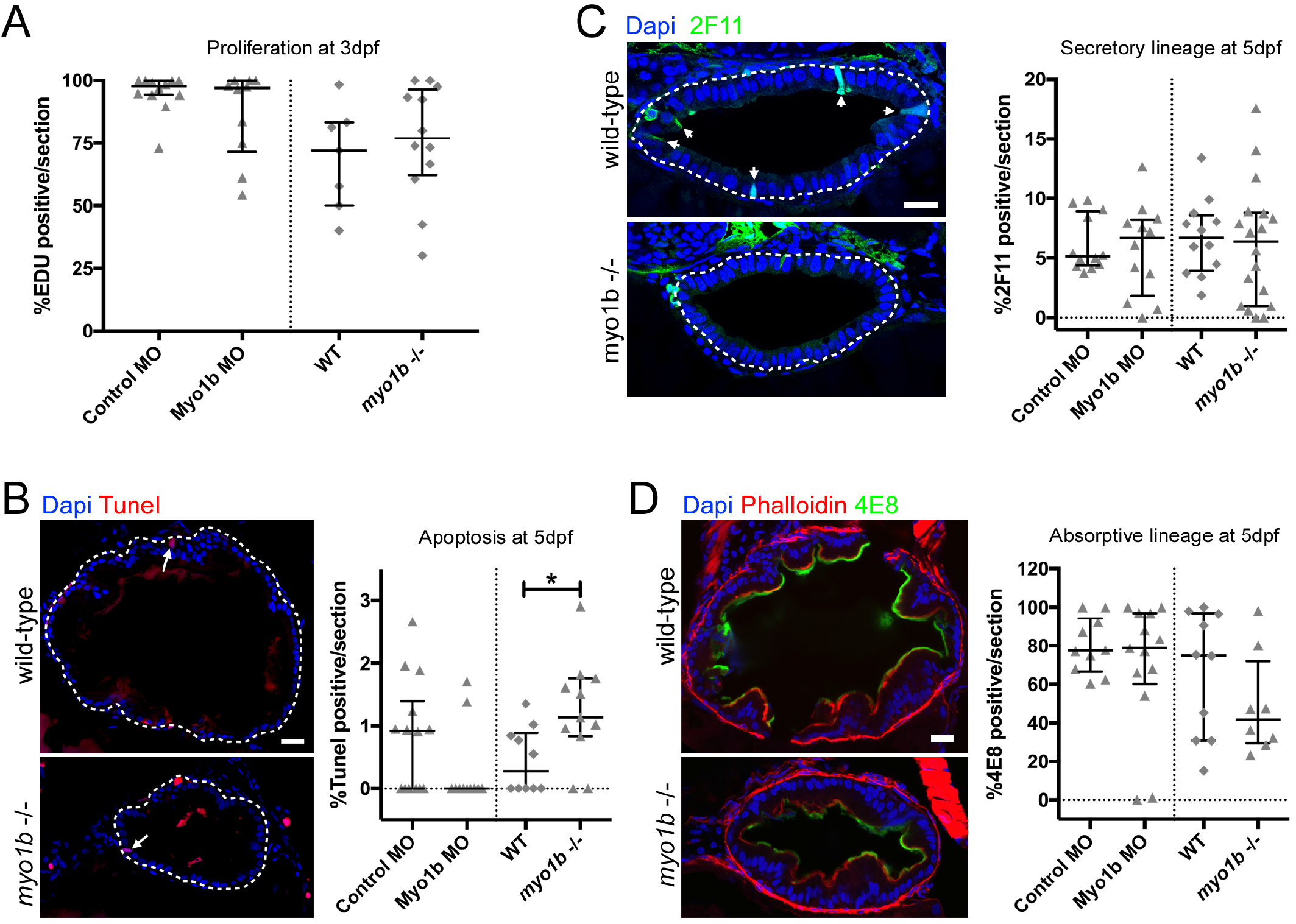
Proliferation, apoptosis and differentiation are essentially unaffected in Myo1b MO and mutant conditions. **A-** Quantifications from EDU and Dapi stained sections of the proportion of cells in S-phase at 3dpf do not reveal significant differences in the proliferative rate of Control (n=12) vs Myo1b MO (n=10) and of WT (n=7) vs *myo1b*-/- (n=12) samples. **B-** Confocal sections of intestinal bulbs stained for apoptosis (Tunel, red) at 5dpf in WT and *myo1b*-/-samples and quantifications of the proportion of apoptotic cells in the four conditions (Control n=14, Myo1b MO n=11; WT n=10; *myo1b*-/-n=11). **C, D-** Confocal sections of intestinal bulbs stained for differentiation markers in WT and *myo1b*-/-samples and quantifications of the proportion of differentiated cells in the four conditions. At 5dpf, neither differentiation of the secretory lineage (**C** - 2F11, Control n=12, Myo1b MO n=12; WT n=12; *myo1b*-/-n=18) nor differentiation of the absorptive lineage (**D -** 4E8, Control n=10, Myo1b MO n=13; WT n=10; *myo1b*-/-n=8) are significantly altered. For all quantifications, data represented are median and interquartile range, Wilcoxon test, *p<0.05. For confocal images, nuclei are counterstained with Dapi (blue), bars=20µm.

**Table S1.**
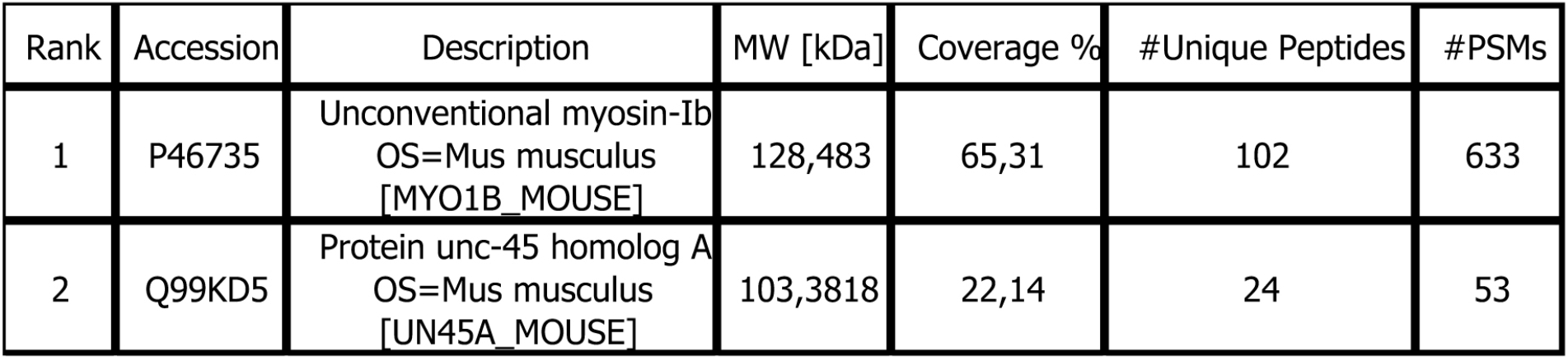
Mass spectrometry result of the GFP-Myo1b pull down assay. UNC45A ranks second, directly after Myo1b. MW, molecular weight; #Unique Peptides, number of distinct peptide sequences identified; # PSMs (peptide spectrum matches) total number of identified peptide sequences for the protein.

| Rank | Accession | Description | MW [kDa] | Coverage % | #Unique Peptides | #PSMs |
| --- | --- | --- | --- | --- | --- | --- |
| 1 | P46735 | Unconventional myosin-Ib<br>OS=Mus musculus<br>[MYO1B_MOUSE] | 128,483 | 65,31 | 102 | 633 |
| 2 | Q99KD5 | Protein unc-45 homolog A<br>OS=Mus musculus<br>[UN45A_MOUSE] | 103,3818 | 22,14 | 24 | 53 |

